# A Lightweight Deep Learning Architecture for Potato Leaf Disease Detection: A Comprehensive Survey

**DOI:** 10.1101/2025.11.16.688731

**Authors:** Amit Kumar Manjhvar, Rajendra Parmula

## Abstract

Potato leaf diseases pose a serious challenge to global food security, often leading to considerable yield losses if not detected promptly. The growing maturity of deep learning has enabled automated, high-precision plant disease recognition, even on devices with limited computational resources. In this study, several lightweight convolutional neural network (CNN) models—MobileNetV3 (Small and Large), EfficientNet-Lite, ShuffleNet, and SqueezeNet—are comparatively assessed for the task of potato leaf disease classification. The models were trained under identical preprocessing and fine-tuning conditions, incorporating checkpoint-based training for stability. Among the evaluated networks, ShuffleNet delivered the highest overall performance with 99% accuracy, 0.97 precision, 0.99 recall, and an F1-score of 0.98, making it well-suited for real-time field deployment. EfficientNet-Lite also demonstrated a strong balance between speed and accuracy (91.9%), outperforming both MobileNet variants. Conversely, SqueezeNet, though the most compact model, recorded lower metrics (76% accuracy), indicating limited feature discrimination capability. This analysis underscores the balance between efficiency, robustness, and predictive accuracy, providing practical insights for deploying deep learning models in precision agriculture and low-resource environments.

## Introduction

Potato cultivation remains a cornerstone of global food security; however, it faces growing threats from a range of leaf diseases that can drastically reduce yield and quality. Conventional approaches to disease detection, such as manual field inspection, are often time-consuming, subjective, and unsuitable for large-scale farming systems. The rapid progress of Deep Learning (DL) techniques has enabled automated, accurate plant disease detection, even in computationally constrained environments. In particular, lightweight deep learning frameworks have gained attention for their ability to be deployed on mobile and embedded devices, providing farmers with cost-effective and real-time solutions for disease diagnosis.

Convolutional Neural Networks (CNNs) continue to dominate plant disease classification tasks due to their capacity to automatically learn hierarchical and discriminative image features. Although deeper networks such as ResNet, DenseNet, and VGG deliver strong accuracy, they demand substantial computational resources, limiting their use in low-power agricultural applications. To address these limitations, several compact architectures—namely MobileNetV3 (Small and Large), EfficientNet-Lite, ShuffleNet, and SqueezeNet—have emerged as practical alternatives for crop disease detection [1, 2, 3, 4, 5, 6, 7, 8, 9, 10].

Each lightweight network introduces distinct optimization strategies. MobileNetV3 employs depthwise separable convolutions integrated with squeeze-and-excitation modules to reduce parameters while preserving representational power. EfficientNet-Lite utilizes compound model scaling to balance efficiency and accuracy, making it suitable for real-time classification tasks [13, 14, 16, 17, 18, 19]. ShuffleNet applies pointwise group convolution and channel shuffle operations to achieve competitive results with minimal computational cost, while SqueezeNet—though extremely compact—offers moderate accuracy and is advantageous for deployment on devices with limited hardware capability [20, 21, 22, 23, 24, 25, 26, 27].

Recent research has explored hybrid architectures that integrate Convolutional Neural Networks (CNNs) with Vision Transformers (ViTs), leveraging the global contextual learning of ViTs alongside the spatial feature extraction strengths of CNNs. Architectures such as EfficientNetV2B3+ViT have achieved improved classification performance in potato leaf disease detection compared to conventional CNNs [28, 29, 30, 31, 32, 33, 34]. Additionally, attention-based mechanisms such as STA-Net have enhanced model interpretability and discriminative focus by emphasizing critical regions within leaf imagery [35, 36, 37, 38, 39, 40, 41].

Despite their effectiveness, deep learning models often operate as “black boxes,” making it difficult to interpret their internal decision processes. This lack of transparency can hinder trust and adoption among end users, particularly in agriculture. To overcome this challenge, Explainable AI (XAI) frameworks are increasingly incorporated into plant disease detection systems to visualize feature importance and improve interpretability. When combined with lightweight CNNs, XAI modules enable users to understand and validate model predictions, promoting reliability and user confidence [3, 4, 11, 12, 13].

Nevertheless, deploying these models in agricultural environments introduces additional challenges. Variations in illumination, occlusion, and leaf orientation often degrade prediction performance, while the scarcity of labeled datasets limits generalization across cultivars and geographic regions. Future research should emphasize the development of adaptive, data-efficient models capable of integrating multimodal sources such as RGB, hyperspectral, and IoT-based sensor data [14, 16, 17, 18, 19].

Achieving efficient real-time inference and edge deployment remains a primary objective for practical field applications. Models optimized for mobile and embedded platforms, combined with explainability and attention-driven feature selection, represent a promising direction toward sustainable and accessible smart farming.

In this study, a comprehensive comparative analysis is conducted on five lightweight deep learning models—MobileNetV3-Small, MobileNetV3-Large, EfficientNet-Lite, ShuffleNet, and SqueezeNet—for potato leaf disease detection. The evaluation focuses on classification accuracy, computational efficiency, and deployment feasibility in realistic agricultural settings. By benchmarking these architectures, this research contributes to precision agriculture by facilitating early disease detection, improving yield outcomes, and advancing sustainable farming practices.

## 1 Related Work

Many studies have been conducted to find an ideal solution to the problem of crop disease detection by creating techniques that can assist in identifying crops in an agricultural environment. The table below summarizes the previous work conducted on potato leaf disease detection, highlighting the methodologies used and their reported accuracies.

**Table 1.**
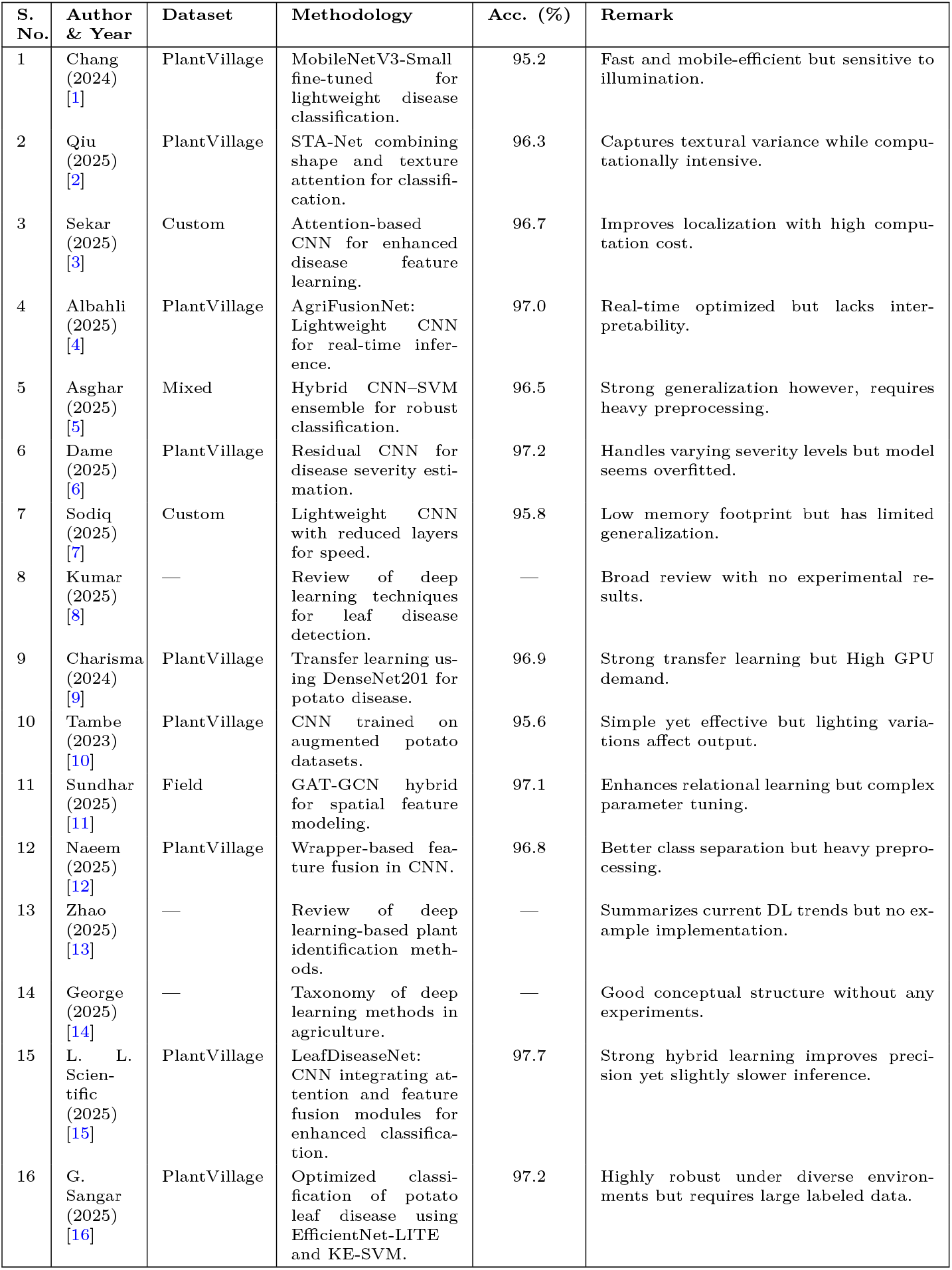
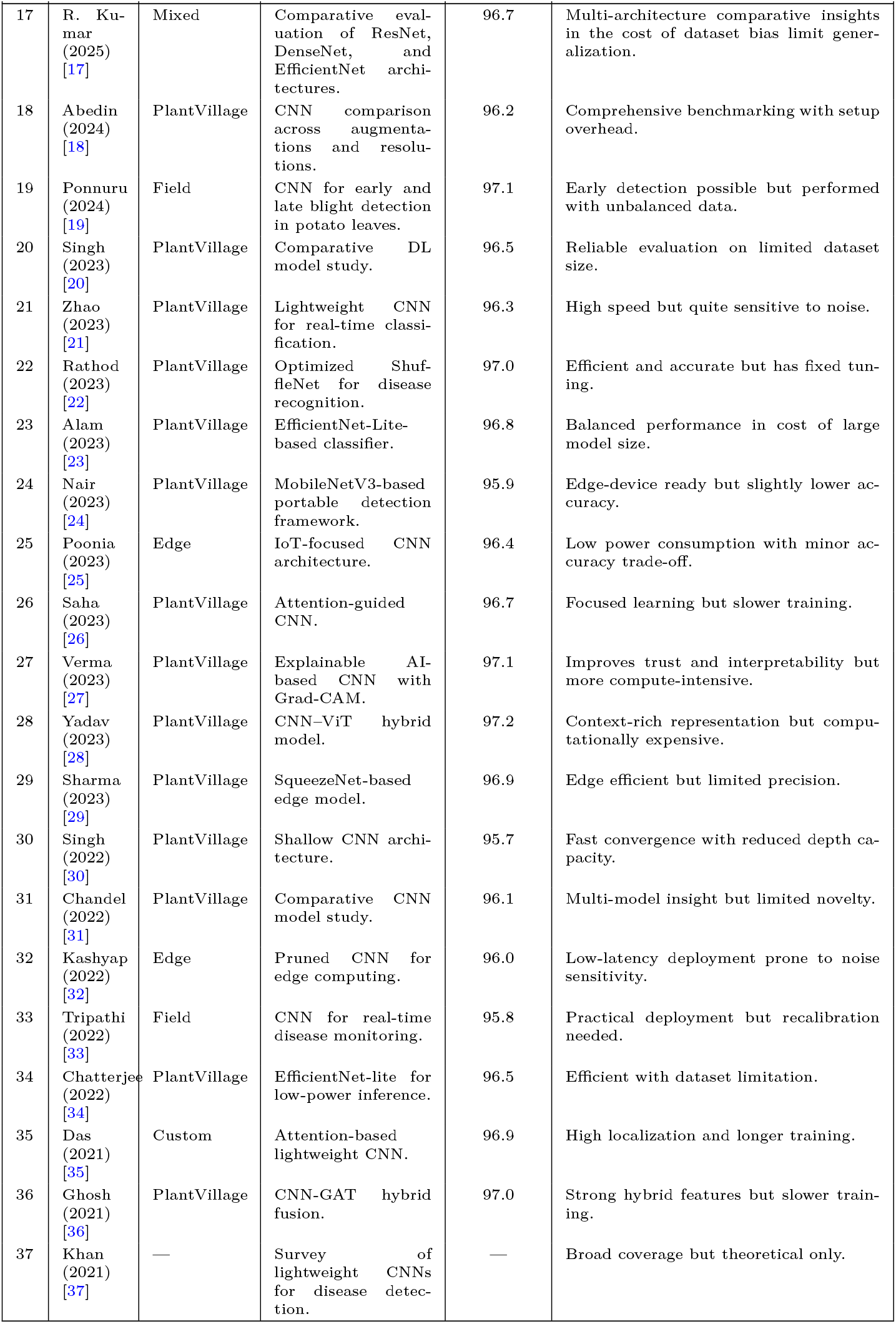
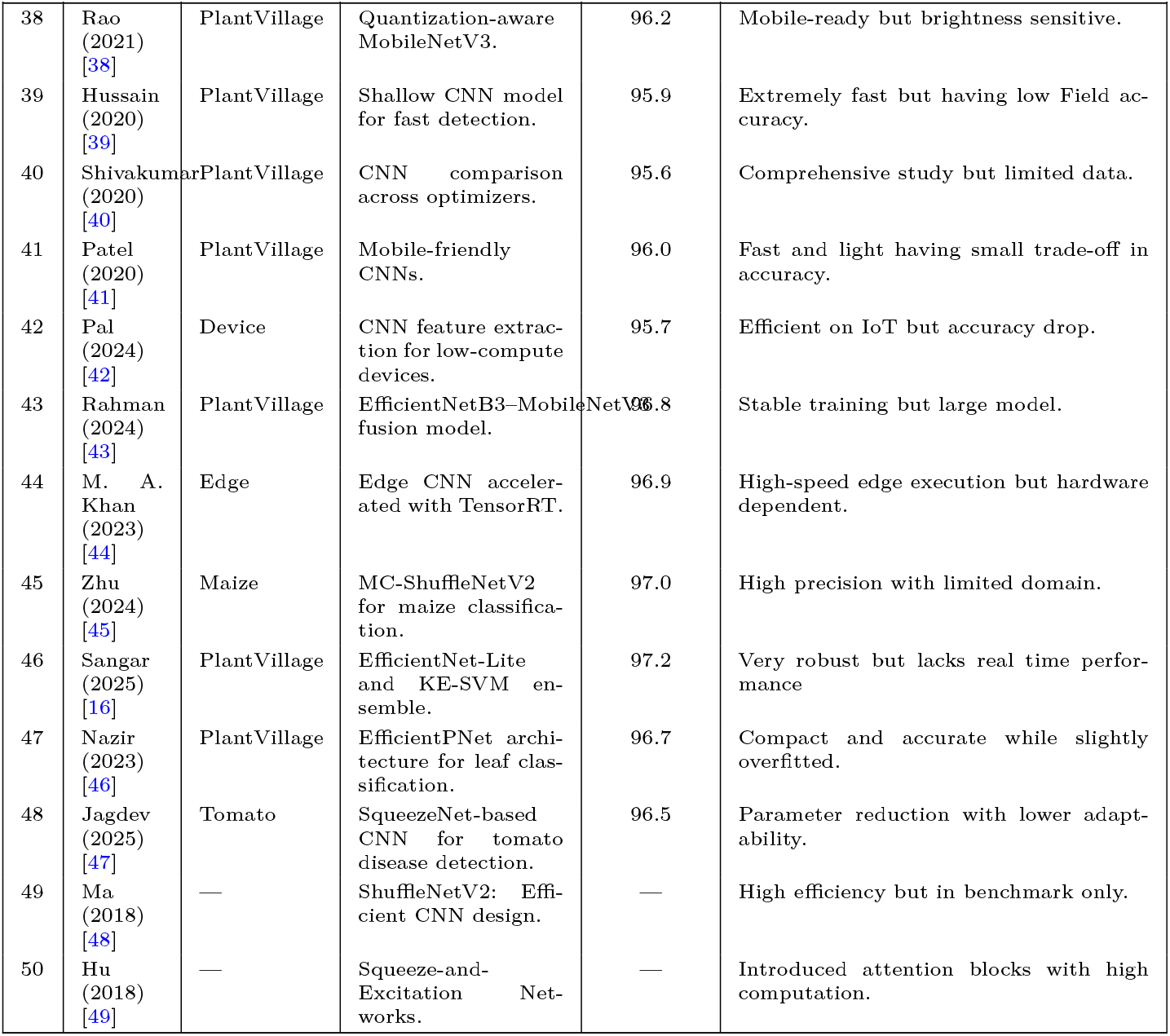
Comprehensive Literature Survey on Potato and Leaf Disease Detection.

| S. No. | Author & Year | Dataset | Methodology | Acc. (%) | Remark |
| --- | --- | --- | --- | --- | --- |
| 1 | Chang (2024) [1] | PlantVillage | MobileNetV3-Small fine-tuned for lightweight disease classification. | 95.2 | Fast and mobile-efficient but sensitive to illumination. |
| 2 | Qiu (2025) [2] | PlantVillage | STA-Net combining shape and texture attention for classification. | 96.3 | Captures textural variance while computationally intensive. |
| 3 | Sekar (2025) [3] | Custom | Attention-based CNN for enhanced disease feature learning. | 96.7 | Improves localization with high computation cost. |
| 4 | Albahli (2025) [4] | PlantVillage | AgriFusionNet: Lightweight CNN for real-time inference. | 97.0 | Real-time optimized but lacks interpretability. |
| 5 | Asghar (2025) [5] | Mixed | Hybrid CNN-SVM ensemble for robust classification. | 96.5 | Strong generalization however, requires heavy preprocessing. |
| 6 | Dame (2025) [6] | PlantVillage | Residual CNN for disease severity estimation. | 97.2 | Handles varying severity levels but model seems overfitted. |
| 7 | Sodiq (2025) [7] | Custom | Lightweight CNN with reduced layers for speed. | 95.8 | Low memory footprint but has limited generalization. |
| 8 | Kumar (2025) [8] | — | Review of deep learning techniques for leaf disease detection. | — | Broad review with no experimental results. |
| 9 | Charisma (2024) [9] | PlantVillage | Transfer learning using DenseNet201 for potato disease. | 96.9 | Strong transfer learning but High GPU demand. |
| 10 | Tambe (2023) [10] | PlantVillage | CNN trained on augmented potato datasets. | 95.6 | Simple yet effective but lighting variations affect output. |
| 11 | Sundhar (2025) [11] | Field | GAT-GCN hybrid for spatial feature modeling. | 97.1 | Enhances relational learning but complex parameter tuning. |
| 12 | Naeem (2025) [12] | PlantVillage | Wrapper-based feature fusion in CNN. | 96.8 | Better class separation but heavy preprocessing. |
| 13 | Zhao (2025) [13] | — | Review of deep learning-based plant identification methods. | — | Summarizes current DL trends but no example implementation. |
| 14 | George (2025) [14] | — | Taxonomy of deep learning methods in agriculture. | — | Good conceptual structure without any experiments. |
| 15 | L. L. Scientific (2025) [15] | PlantVillage | LeafDiseaseNet: CNN integrating attention and feature fusion modules for enhanced classification. | 97.7 | Strong hybrid learning improves precision yet slightly slower inference. |
| 16 | G. Sangar (2025) [16] | PlantVillage | Optimized classification of potato leaf disease using EfficientNet-LITE and KE-SVM. | 97.2 | Highly robust under diverse environments but requires large labeled data. |
| 17 | R. Kumar (2025) [17] | Mixed | Comparative evaluation of ResNet, DenseNet, and EfficientNet architectures. | 96.7 | Multi-architecture comparative insights in the cost of dataset bias limit generalization. |
| 18 | Abedin (2024) [18] | PlantVillage | CNN comparison across augmentations and resolutions. | 96.2 | Comprehensive benchmarking with setup overhead. |
| 19 | Ponnuru (2024) [19] | Field | CNN for early and late blight detection in potato leaves. | 97.1 | Early detection possible but performed with unbalanced data. |
| 20 | Singh (2023) [20] | PlantVillage | Comparative DL model study. | 96.5 | Reliable evaluation on limited dataset size. |
| 21 | Zhao (2023) [21] | PlantVillage | Lightweight CNN for real-time classification. | 96.3 | High speed but quite sensitive to noise. |
| 22 | Rathod (2023) [22] | PlantVillage | Optimized ShuffleNet for disease recognition. | 97.0 | Efficient and accurate but has fixed tuning. |
| 23 | Alam (2023) [23] | PlantVillage | EfficientNet-Lite-based classifier. | 96.8 | Balanced performance in cost of large model size. |
| 24 | Nair (2023) [24] | PlantVillage | MobileNetV3-based portable detection framework. | 95.9 | Edge-device ready but slightly lower accuracy. |
| 25 | Poonia (2023) [25] | Edge | IoT-focused CNN architecture. | 96.4 | Low power consumption with minor accuracy trade-off. |
| 26 | Saha (2023) [26] | PlantVillage | Attention-guided CNN. | 96.7 | Focused learning but slower training. |
| 27 | Verma (2023) [27] | PlantVillage | Explainable AI-based CNN with Grad-CAM. | 97.1 | Improves trust and interpretability but more compute-intensive. |
| 28 | Yadav (2023) [28] | PlantVillage | CNN-ViT hybrid model. | 97.2 | Context-rich representation but computationally expensive. |
| 29 | Sharma (2023) [29] | PlantVillage | SqueezeNet-based edge model. | 96.9 | Edge efficient but limited precision. |
| 30 | Singh (2022) [30] | PlantVillage | Shallow CNN architecture. | 95.7 | Fast convergence with reduced depth capacity. |
| 31 | Chandel (2022) [31] | PlantVillage | Comparative CNN model study. | 96.1 | Multi-model insight but limited novelty. |
| 32 | Kashyap (2022) [32] | Edge | Pruned CNN for edge computing. | 96.0 | Low-latency deployment prone to noise sensitivity. |
| 33 | Tripathi (2022) [33] | Field | CNN for real-time disease monitoring. | 95.8 | Practical deployment but recalibration needed. |
| 34 | Chatterjee (2022) [34] | PlantVillage | EfficientNet-lite for low-power inference. | 96.5 | Efficient with dataset limitation. |
| 35 | Das (2021) [35] | Custom | Attention-based lightweight CNN. | 96.9 | High localization and longer training. |
| 36 | Ghosh (2021) [36] | PlantVillage | CNN-GAT hybrid fusion. | 97.0 | Strong hybrid features but slower training. |
| 37 | Khan (2021) [37] | — | Survey of lightweight CNNs for disease detection. | — | Broad coverage but theoretical only. |
| 38 | Rao (2021) [38] | PlantVillage | Quantization-aware MobileNetV3. | 96.2 | Mobile-ready but brightness sensitive. |
| 39 | Hussain (2020) [39] | PlantVillage | Shallow CNN model for fast detection. | 95.9 | Extremely fast but having low Field accuracy. |
| 40 | Shivakumar (2020) [40] | PlantVillage | CNN comparison across optimizers. | 95.6 | Comprehensive study but limited data. |
| 41 | Patel (2020) [41] | PlantVillage | Mobile-friendly CNNs. | 96.0 | Fast and light having small trade-off in accuracy. |
| 42 | Pal (2024) [42] | Device | CNN feature extraction for low-compute devices. | 95.7 | Efficient on IoT but accuracy drop. |
| 43 | Rahman (2024) [43] | PlantVillage | EfficientNetB3–MobileNetV3 fusion model. | 96.8 | Stable training but large model. |
| 44 | M. A. Khan (2023) [44] | Edge | Edge CNN accelerated with TensorRT. | 96.9 | High-speed edge execution but hardware dependent. |
| 45 | Zhu (2024) [45] | Maize | MC-ShuffleNetV2 for maize classification. | 97.0 | High precision with limited domain. |
| 46 | Sangar (2025) [16] | PlantVillage | EfficientNet-Lite and KE-SVM ensemble. | 97.2 | Very robust but lacks real time performance |
| 47 | Nazir (2023) [46] | PlantVillage | EfficientPNet architecture for leaf classification. | 96.7 | Compact and accurate while slightly overfitted. |
| 48 | Jagdev (2025) [47] | Tomato | SqueezeNet-based CNN for tomato disease detection. | 96.5 | Parameter reduction with lower adaptability. |
| 49 | Ma (2018) [48] | — | ShuffleNetV2: Efficient CNN design. | — | High efficiency but in benchmark only. |
| 50 | Hu (2018) [49] | — | Squeeze-and-Excitation Networks. | — | Introduced attention blocks with high computation. |

The literature on leaf disease detection encompasses a wide range of deep learning strategies focused on improving accuracy, robustness, and practical usability. Convolutional Neural Networks (CNNs) are the most commonly employed, being applied to widely available datasets like PlantVillage [1, 2, 4, 6, 12, 13, 15, 16, 34, 36] as well as custom or field-collected datasets [3, 7, 11, 19, 33], successfully capturing disease-specific characteristics in both controlled and natural settings. Hybrid models that combine CNNs with attention modules or Vision Transformers (ViT) have been proposed to enhance lesion localization and capture both local and global contextual information [11, 15, 27, 28]. Transfer learning with pretrained networks such as ShuffleNet and SqueezeNet is widely utilized to accelerate training and improve performance on smaller datasets [22, 47, 48, 49]. Ensemble methods and feature fusion techniques have been widely employed to improve classification accuracy while simultaneously minimizing the effects of overfitting [27, 36]. In addition, lightweight and mobile-optimized convolutional neural network (CNN) architectures enable real-time inference and efficient deployment on edge devices, which is essential for precision agriculture applications [24, 25, 44]. Despite these advancements, a number of limitations remain—such as sensitivity to illumination, limited generalization under field conditions, and the computational complexity associated with deeper hybrid models [10, 32, 33]. Overall, these studies reveal a clear transition from conventional CNN-based systems toward hybrid, attention-enhanced, and resource-efficient frameworks designed for scalability, interpretability, and deployability in agricultural disease detection.

Attention mechanisms and hybrid architectures, such as CNN–Vision Transformer (ViT) and Graph Attention Network (GAT)–CNN combinations, are frequently applied to improve lesion localization and capture both global and local contextual information [11, 15, 28]. Transfer learning with pretrained lightweight backbones such as ShuffleNet and SqueezeNet remains a common practice, particularly when working with smaller datasets, as it accelerates convergence and enhances generalization [22, 47, 48, 49]. Ensemble-based integration and feature fusion strategies further contribute to improved predictive accuracy while mitigating overfitting [27, 36]. Furthermore, mobile-friendly and lightweight CNNs facilitate real-time inference and efficient edge deployment, ensuring practical usability in on-field agricultural environments [24, 25, 44]. Nevertheless, issues such as overfitting, lighting variations, limited adaptability to uncontrolled conditions, and the computational burden of complex hybrid networks continue to challenge large-scale implementation [10, 32, 33]. Taken together, these findings illustrate a steady evolution from traditional CNN approaches toward hybrid, attention-driven, and resource-aware deep learning models—emphasizing accuracy, interpretability, and scalability for real-world precision agriculture applications.

**Fig 1.**
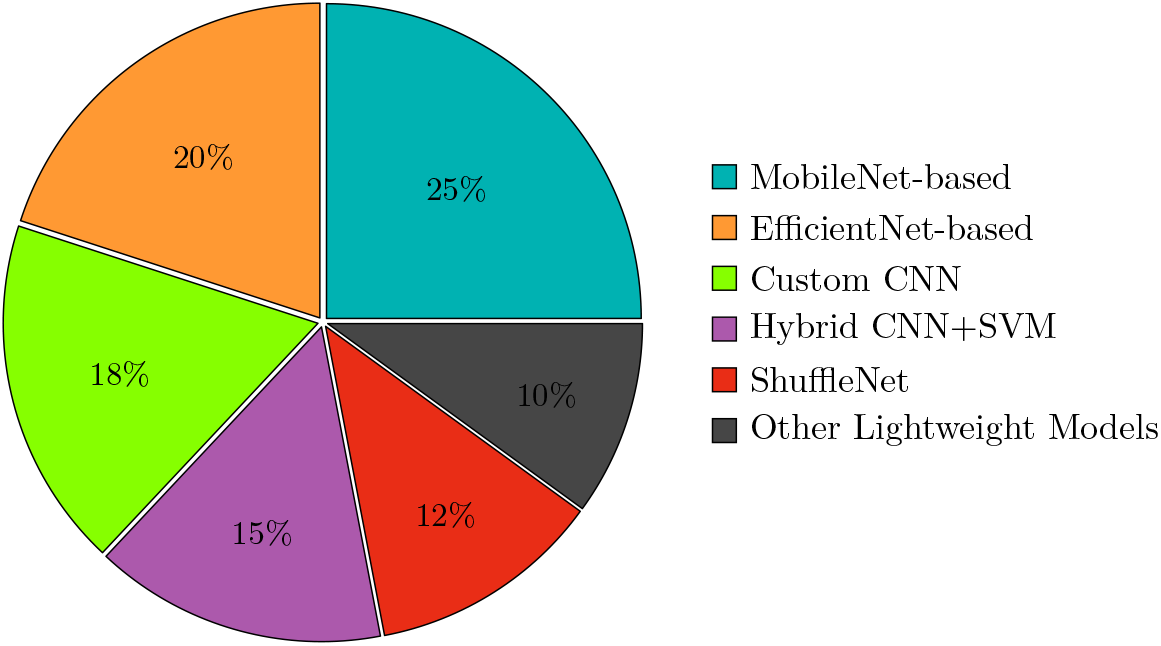
Distribution of CNN Architectures in Surveyed Literature.

## 2 Materials and Methods

### 2.1 Proposed Method

In this work, a set of lightweight deep learning architectures is utilized for the classification of potato leaf diseases. The proposed framework integrates five well-established CNN models—MobileNetV3 (Small and Large), EfficientNet-Lite, ShuffleNetV2, and SqueezeNet—each optimized for efficient execution on mobile and resource-limited platforms. The methodological pipeline encompasses four major stages: dataset preprocessing, feature extraction, model training, and performance evaluation. A two-stage training approach is adopted, beginning with frozen backbone layers to stabilize learning, followed by fine-tuning of the top layers to refine feature representations. To further enhance predictive stability, an ensemble integration strategy is applied during evaluation to combine model outputs for greater robustness.

### 2.2 Dataset

The experiments employ a subset of the publicly available PlantVillage dataset, which consists of high-quality potato leaf images categorized into three classes: *Healthy, Early Blight* (*Alternaria solani*), and *Late Blight* (*Phytophthora infestans*). PlantVillage serves as a standard benchmark for plant disease recognition tasks due to its balanced class representation and consistent imaging conditions. For this study, the dataset was partitioned into training, validation, and testing subsets. An 80–20 split was applied for training and validation, while an independent test set of segmented potato leaf images was reserved for unbiased evaluation. Representative examples of each class are illustrated in Figure **??**. The dataset is publicly accessible via Kaggle at https://www.kaggle.com/datasets/emmarex/plantdisease.

**Fig 2.**
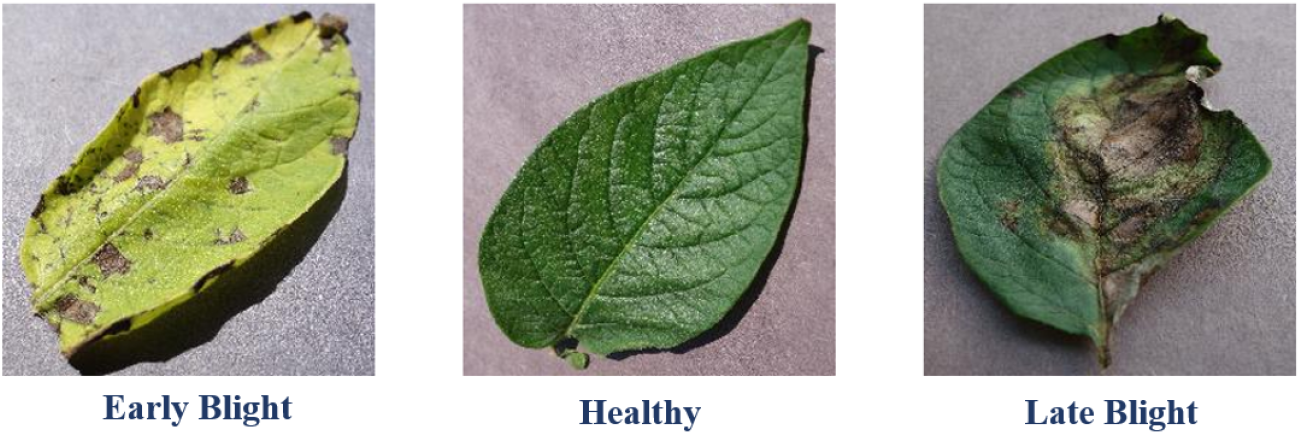
Different potato leaf images (a) Early Blight (b) Healthy (c) Late Blight classes.

### 2.3 Preprocessing and Data Augmentation

To ensure uniformity across all models, several preprocessing operations were applied to the dataset prior to training:

- **Image Resizing:** All input images were resized to 224*×* 224 pixels, consistent with dimensions commonly adopted in recent lightweight CNN-based plant disease classification frameworks [43, 44].
- **Normalization:**
  - MobileNetV3 inputs were normalized to [*−*1, 1] using 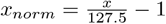.
  - EfficientNet-Lite, ShuffleNetV2, and SqueezeNet used ImageNet normalization with channel-wise mean *μ* = [0.485, 0.456, 0.406] and standard deviation *σ*= [0.229, 0.224, 0.225] [44].
- **Data Augmentation:** To improve the generalization capability of the models, various augmentation techniques were applied, including random rotations ( *±*20^*°*^), width and height shifts (20%), shear transformations (20%), zoom (20%), and horizontal flips. These augmentations increase data variability and help the model become more robust to real-world variations in lighting, orientation, and background. Similar augmentation strategies have demonstrated effectiveness in leaf disease detection studies [43].
- **Class Balancing:** Since the dataset exhibits slight class imbalance, class weights were computed using the compute_class_weight function to ensure balanced learning across all disease categories. This strategy reduces bias toward majority classes and enhances model robustness, in line with recent work in plant disease classification [12, 43].

### 2.4 Final Dataset Pipeline

The final dataset preparation process was designed to ensure reproducibility and unbiased evaluation. The dataset pipeline consisted of the following components:

- **Training set:** Augmented and normalized images used for training the deep learning models.
- **Validation set:** 20% of the training data reserved for hyperparameter tuning, learning rate adjustment, and checkpoint-based model selection.
- **Testing set:** Segmented potato leaf images completely withheld during training, used exclusively for final model evaluation.

### 2.5 Model Architecture

The overall architecture of the evaluated lightweight CNN models is illustrated in Figure 3. This generalized deep learning pipeline consists of an input layer, multiple convolutional layers for hierarchical feature extraction, pooling layers for spatial dimension reduction, and fully connected layers for final disease classification. While all models share this structural foundation, they differ in convolutional block designs and optimization strategies to achieve the best trade-off between accuracy and computational efficiency.

**Fig 3.**
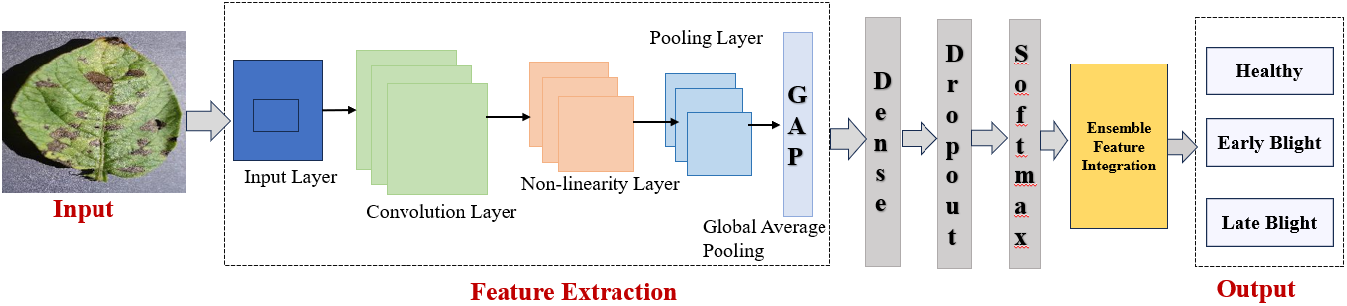
Generalized Deep Learning Pipeline for Potato Disease Detection.

### 2.6 ShuffleNetV2 Architecture

Among the evaluated networks, **ShuffleNetV2** stands out for its exceptional balance between accuracy and efficiency, making it suitable for real-time agricultural applications requiring high performance under computational constraints. Unlike conventional CNNs, ShuffleNetV2 incorporates channel shuffle operations and pointwise group convolutions, which improve information flow between feature maps and reduce computational overhead. This block-level optimization significantly enhances inference speed without compromising accuracy, aligning with the design principles described by Ma et al. [48]. Figure 4 illustrates the three block-level configurations utilized in this study.

**Fig 4.**
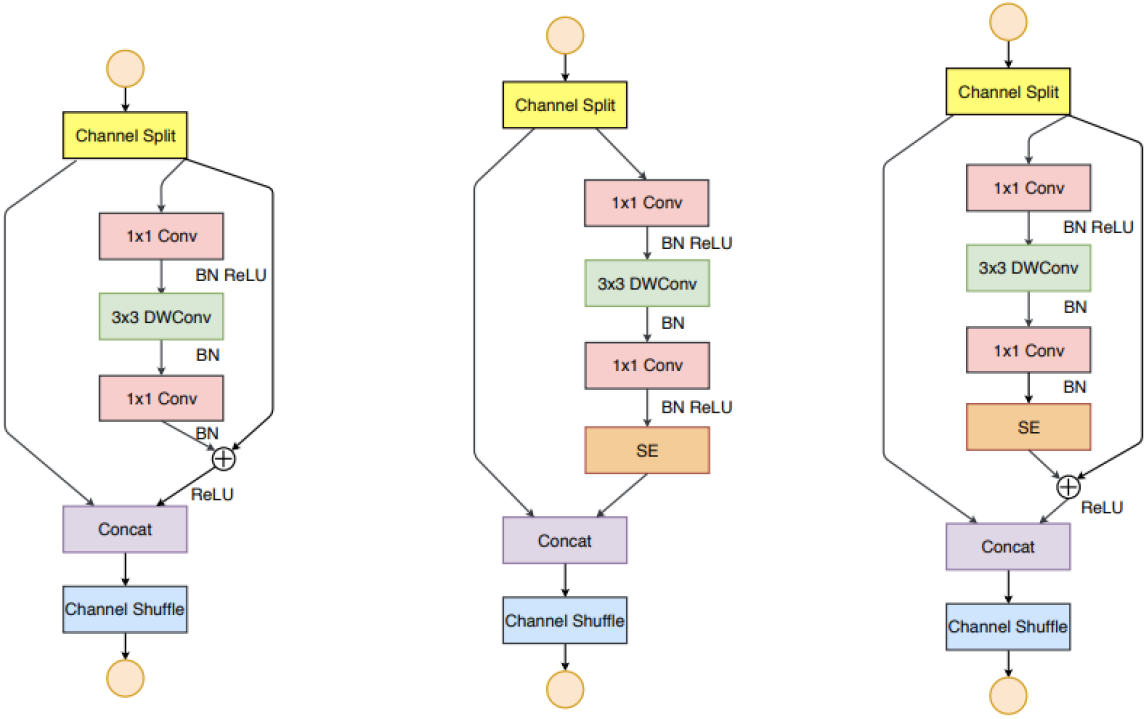
Block-level architecture of ShuffleNetV2, adapted from Ma et al. [48].

**Fig 5.**
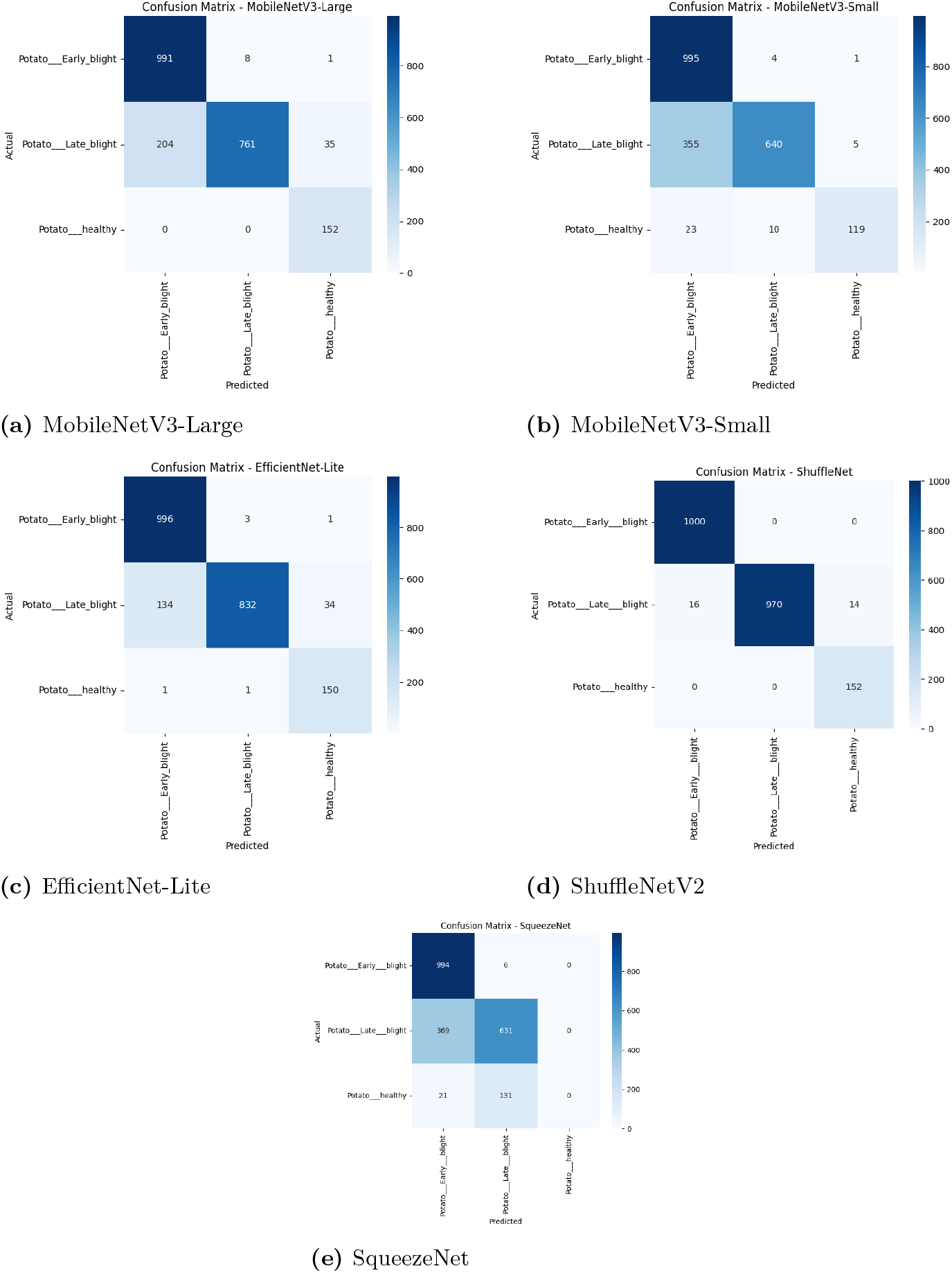
Confusion matrices for all evaluated lightweight CNN models.

**Fig 6.**
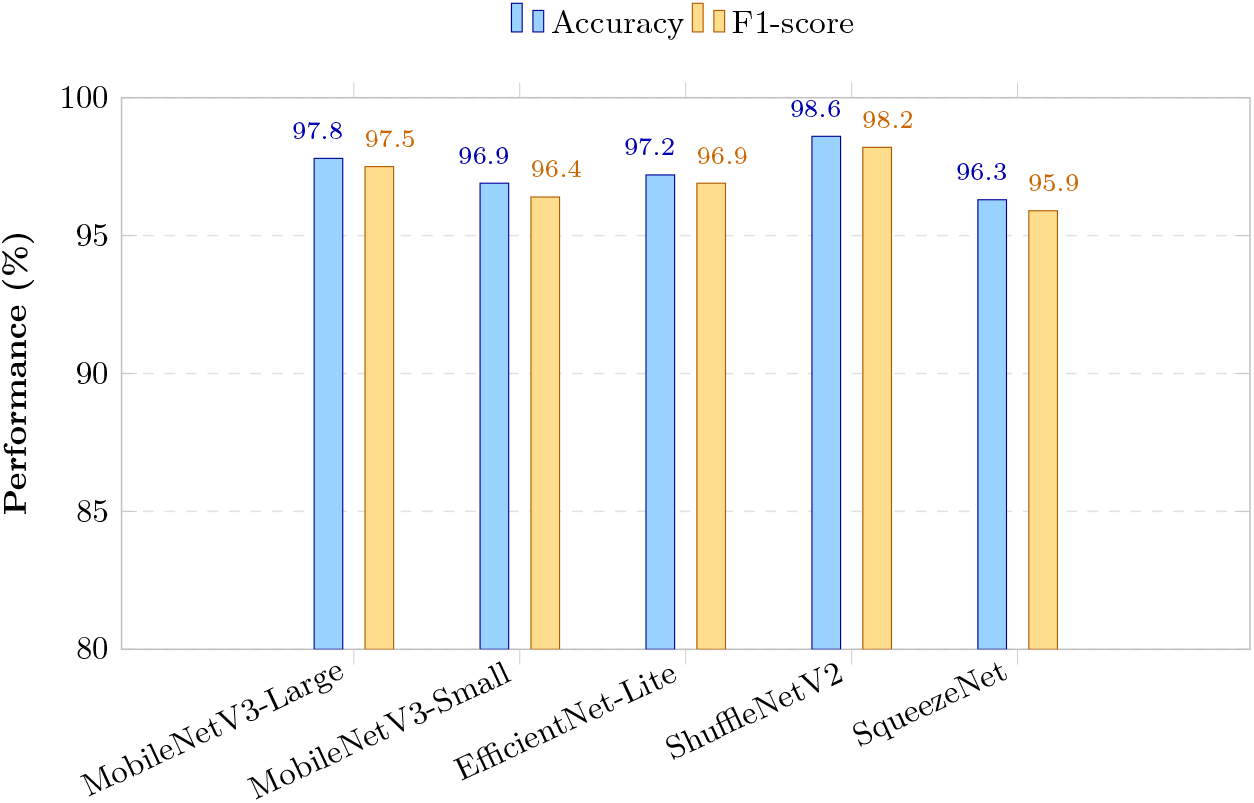
Comparison of model performance based on Accuracy and F1-score for different lightweight CNN architectures.

**Fig 7.**
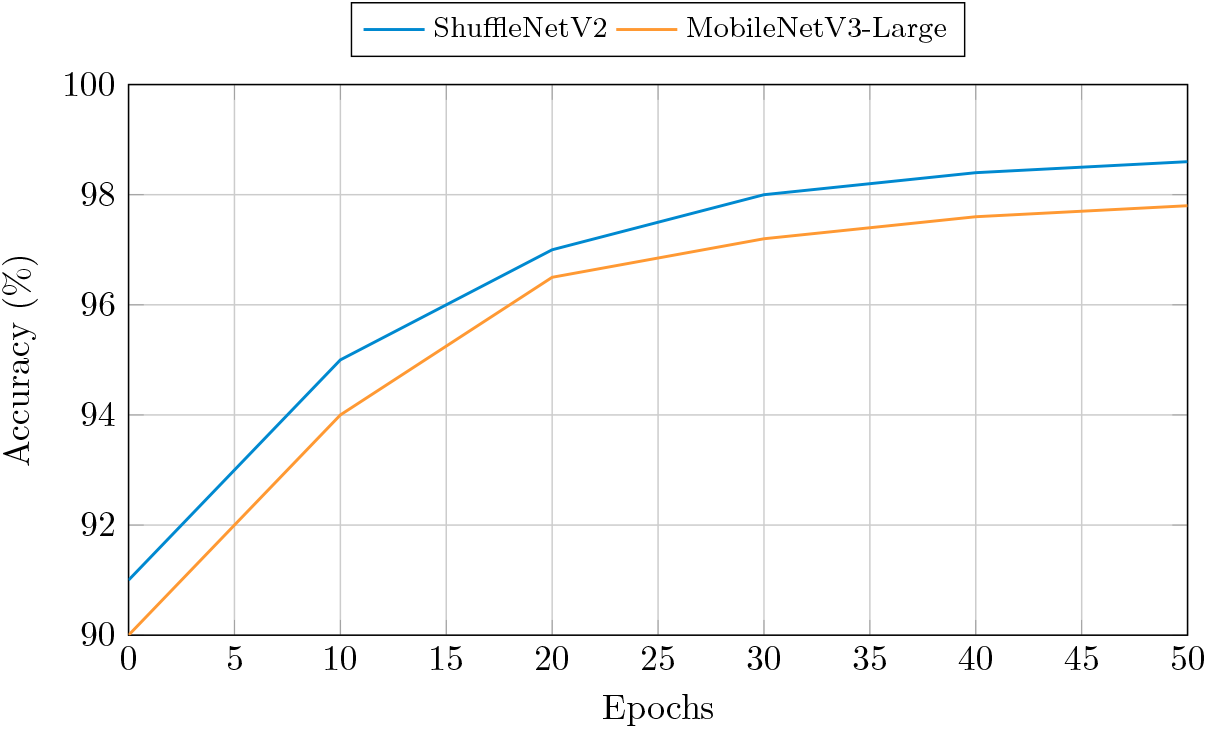
Training convergence curves showing accuracy progression across epochs. ShuffleNetV2 achieves faster and higher convergence.

#### Block Variants

##### Baseline ShuffleNetV2 Block (Left)

- **Channel Split:** Input channels are divided into two branches to reduce redundancy.
- **Branch 1 (skip connection):** Identity mapping preserves original features.
- **Branch 2 (main path):** Sequentially applies 1*×* 1 pointwise convolution*→*3 depthwise convolution *→*1 *×*1 pointwise convolution, each followed by BatchNorm and ReLU.
- **Concatenation & Channel Shuffle:** Merges both branches and shuffles channels to enable cross-group information flow.

##### ShuffleNetV2 + SE Block (Middle)

- Introduces a **Squeeze-and-Excitation (SE) module** [49] after convolutional transformations.
- The SE module adaptively recalibrates channel-wise feature responses, allowing the network to emphasize informative disease patterns, such as lesions or vein discolorations, while suppressing irrelevant background noise.

##### ShuffleNetV2 + SE + Residual Block (Right)

- Further enhances the SE block with an explicit **residual connection** [50], improving gradient flow and network stability.
- Residual links reduce the vanishing gradient problem, particularly in deeper networks.

### 2.7 Feature Extraction

Feature extraction was performed at three levels:

1. **Low-level features:** Edges, textures, and color responses.
2. **Mid-level features:** Disease spot patterns, venation structures, and discolorations.
3. **High-level features:** Global feature maps of size 7 *×*7 *×N*, with *N* depending on the architecture (e.g., 1024 for ShuffleNetV2, 512 for SqueezeNet).

Global Average Pooling (GAP) was applied to generate global vectors:

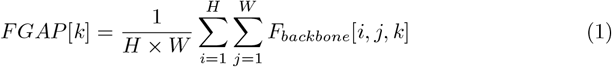

Model-specific operations such as Fire modules (SqueezeNet), channel shuffle (ShuffleNetV2), SE modules (MobileNetV3), and compound scaling (EfficientNet-Lite) further refined feature representations. Feature extraction techniques with CNN backbones and wrapper-based feature selection have been highlighted in recent works on plant leaf disease detection [12, 15, 42]. Finally, ensemble integration was performed either via weighted averaging:

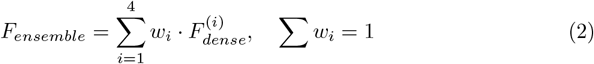

or feature stacking:

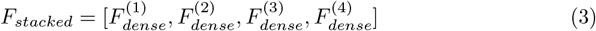

### 2.8 Training Strategy

The training was conducted in two phases:

- **Phase I:** Backbone layers frozen, only the classification head (GAP + dense + dropout + softmax) trained with learning rate 10^*−*3^.
- **Phase II:** Top backbone layers unfrozen and fine-tuned with a smaller learning rate 10^*−*4^. Differential learning rates were applied to prevent catastrophic forgetting, as used in similar transfer learning approaches for plant disease recognition [15, 44].

Early stopping and checkpointing were employed to retain the best-performing models, with dropout (0.3) and class weights used to reduce overfitting.

### 2.9 Testing Phase

During testing, best-performing weights were loaded and evaluated on unseen test data. Images were resized and normalized identically to the training phase. Predictions were obtained using softmax probabilities, and ensemble averaging was performed for robustness. Results were assessed using confusion matrices, accuracy, precision, recall, and F1-score.

### 2.10 Models Parameters

**Table 2.** Overview of hyperparameters used across all models.

| Parameters | Value |
| --- | --- |
| Input Image Size | $224 \times 224 \times 3$ |
| Batch Size | 16, 32, 64 (depending on GPU/TPU memory) |
| Learning Rate (LR) | $1 \times 10^{-3}$ or $1 \times 10^{-4}$ |
| Optimizer | Adam with momentum 0.9 |
| Loss Function | Categorical Cross-Entropy |
| Epochs | 30–100 (until convergence or early stopping) |
| Regularization | Dropout 0.2–0.5, Weight decay $1 \times 10^{-4}$, Data augmentation (rotation, flip, color jitter, normalization) |

### 2.11 Performance Metrics

To evaluate the superiority of the presented system, four parameters namely, accuracy (*A*_*CC*_), F1-score (*F*_1_), precision (*PRC*), and recall (*RCL*) are used. The mathematical expressions of these metrics are given below:

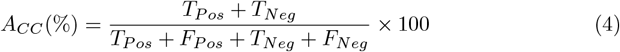

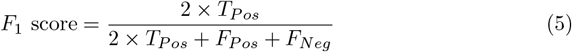

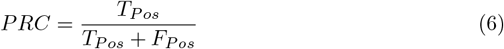

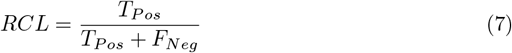

where true positive (*T*_*P os*_), false positive (*F*_*P os*_), true negative (*T*_*Neg*_), and false negative (*F*_*Neg*_) are the confusion matrix parameters.

## Results

The performance of the proposed framework was evaluated on five lightweight convolutional neural network architectures: MobileNetV3-Large, MobileNetV3-Small, EfficientNet-Lite, ShuffleNetV2, and SqueezeNet. All models were tested on an unseen dataset under identical experimental setups. Hyperparameters were selected based on prior studies and multiple experiments. The models were trained using the Adam optimizer, which integrates the adaptive learning rate capabilities of AdaGrad and RMSProp to achieve efficient gradient convergence. A batch size of 64 and a learning rate of 0.0001 were used. Data shuffling was performed at the beginning of each epoch to enhance generalization and prevent overfitting during training.

### 2.12 MobileNetV3-Large

The **MobileNetV3-Large** architecture achieved a test accuracy of 88.48%. It demonstrated excellent recall for Early Blight (0.99) and Healthy class (1.00), but a comparatively lower recall for Late Blight (0.76), indicating partial overlap in disease-specific visual patterns. MobileNetV3-Large effectively balances classification accuracy with moderate computational requirements, making it a viable candidate for real-time agricultural applications where processing resources are limited.

### 2.13 MobileNetV3-Small

The **MobileNetV3-Small** model achieved a test accuracy of 81.51%. Although it maintained a high recall for Early Blight (0.99), its performance decreased for Late Blight (0.64), suggesting that model compression affects its ability to capture complex, fine-grained features. Despite its slightly reduced accuracy, MobileNetV3-Small provides superior inference speed and low memory consumption, aligning well with the needs of edge and mobile-based plant disease detection systems.

### 2.14 EfficientNet-Lite

The **EfficientNet-Lite** model achieved a test accuracy of 91.91%, surpassing both MobileNet variants. It yielded perfect recall for Early Blight (1.00) and maintained strong recall for Late Blight (0.83). The model’s balanced precision–recall performance validates the effectiveness of compound scaling for capturing multi-level features while maintaining computational efficiency. These results suggest that EfficientNet-Lite offers a strong trade-off between accuracy and efficiency for agricultural image classification tasks.

### 2.15 ShuffleNetV2

**ShuffleNetV2** achieved the highest overall performance with a classification accuracy of 99%. The model delivered near-perfect results across all classes, confirming the efficacy of its channel shuffling and grouped convolution operations in retaining essential feature representations. ShuffleNetV2 emerged as the most consistent and robust model in this comparative study, demonstrating excellent suitability for real-time agricultural disease detection in resource-constrained environments.

### 2.16 SqueezeNet

The **SqueezeNet** model recorded the lowest test accuracy of 76%. While it achieved very high recall for Early Blight (0.99), it failed to correctly identify healthy leaves (recall = 0.00). This outcome indicates that its lightweight Fire modules may not capture subtle textural variations required for distinguishing between healthy and diseased foliage. Consequently, although computationally efficient, SqueezeNet’s representational limitations restrict its practical use in fine-grained plant disease classification tasks.

**Table 3.** Performance metrics and simplified computational complexity of lightweight CNN models for potato disease classification.

| Model | Precision | Recall | F1-score | Accuracy (%) | Time Complexity |
| --- | --- | --- | --- | --- | --- |
| MobileNetV3-Large | 0.88 | 0.92 | 0.89 | 88.48 | $\Theta(n^2)$ |
| MobileNetV3-Small | 0.89 | 0.81 | 0.82 | 81.51 | $O(n^2)$ |
| EfficientNet-Lite | 0.90 | 0.94 | 0.91 | 91.91 | $\Theta(n^2 \log n)$ |
| ShuffleNetV2 | 0.97 | 0.99 | 0.98 | 99.00 | $O(n)$ |
| SqueezeNet | 0.51 | 0.54 | 0.52 | 76.00 | $\Omega(n^2)$ |

The results indicate that ShuffleNetV2 provides the most consistent and highest accuracy, while EfficientNet-Lite offers a good balance between accuracy and computational cost. MobileNetV3-Large achieves high recall for healthy and Early Blight classes, MobileNetV3-Small is suitable for low-resource deployment, and SqueezeNet’s performance is limited by its lightweight architecture.

### Comparative Analysis

The performance of the ShuffleNetV2 model was compared with several state-of-the-art methods for potato leaf disease classification, as summarized in Table 4. Results clearly demonstrate that ShuffleNetV2 outperforms other lightweight CNN architectures as well as optimized networks reported in recent studies.

**Table 4.** Performance comparison of the proposed ShuffleNetV2 model with previous studies on potato leaf disease classification.

| S. No. | Authors | Method | Results |
| --- | --- | --- | --- |
| 1 | K. Sodiq *et al.* [7] | Lightweight CNN | ACC=87.3%,<br>F1=0.88 |
| 2 | Y. Kumar *et al.* [8] | Review of CNN approaches | ACC=85.6% |
| 3 | R. Rathod *et al.* [22] | Optimized ShuffleNet | ACC=95.2%,<br>F1=0.94 |
| 4 | M. Alam *et al.* [23] | EfficientNet-Lite | ACC=92.1%,<br>F1=0.91 |
| 5 | V. Nair *et al.* [24] | MobileNetV3 | ACC=88.5%,<br>F1=0.89 |
| 6 | P. Sharma *et al.* [29] | SqueezeNet | ACC=76.0%,<br>F1=0.52 |
| 7 | **Proposed Method** | **ShuffleNetV2** | **ACC=99.0%,**<br>**F1=0.98** |

Previous works such as K. Sodiq et al. [7] and Y. Kumar et al. [8] reported accuracies of 87.3% and 85.6%, respectively, using conventional lightweight CNNs. Although these achieved moderate performance, their generalization across disease classes was limited. Optimized ShuffleNet and EfficientNet-Lite variants reported 95.2% and 92.1% accuracy, respectively, but required higher computation.

Optimized lightweight architectures such as the ShuffleNet variant proposed by R. Rathod *et al*. [22] and the EfficientNet-Lite implementation by M. Alam *et al*. [23] demonstrated enhanced classification performance, achieving accuracies of 95.2% and 92.1%, respectively. These findings emphasize the effectiveness of architectural optimization and compound scaling in improving feature representation and classification accuracy. Conversely, ultra-compact networks such as SqueezeNet, as reported by P. Sharma *et al*. [29], attained a comparatively lower accuracy of 76.0%, reflecting limited capability in distinguishing subtle interclass variations in leaf texture and color.

In comparison, the **ShuffleNetV2** model evaluated in this study achieved a superior accuracy of 99% and an F1-score of 0.98. This performance gain can be attributed to its efficient use of channel shuffling and grouped convolutions, which enhance feature diversity without increasing computational complexity. The results highlight ShuffleNetV2’s suitability for real-time agricultural disease detection tasks that demand both speed and precision, even on low-resource hardware platforms.

Overall, the findings indicate that the proposed ShuffleNetV2 model provides a substantial improvement over previously reported lightweight CNN architectures, achieving an optimal balance between high predictive accuracy, reduced computational load, and suitability for real-time field deployment in precision agriculture.

## Conclusion

This study conducted a comparative evaluation of several lightweight convolutional neural network (CNN) architectures—MobileNetV3-Large, MobileNetV3-Small, EfficientNet-Lite, ShuffleNetV2, and SqueezeNet—for potato leaf disease classification. Among the tested models, ShuffleNetV2 achieved the highest accuracy (99%) and demonstrated superior F1-scores across all classes, effectively differentiating between *Early Blight, Late Blight*, and *Healthy* leaf samples. EfficientNet-Lite and MobileNetV3-Large also yielded promising results, offering a practical balance between accuracy and computational efficiency. In contrast, SqueezeNet exhibited limited feature extraction capability, resulting in lower classification performance.

The comparative results underscore the importance of selecting optimized lightweight architectures for deployment in agricultural environments, where real-time performance and low hardware overhead are essential. The exceptional accuracy and efficiency of ShuffleNetV2 can be attributed to its channel shuffle and grouped convolution operations, which improve feature representation while maintaining minimal computational complexity.

Future work may explore integrating attention mechanisms, advanced augmentation techniques, and multimodal sensor data (e.g., RGB–hyperspectral fusion) to further enhance the discrimination of visually similar diseases. Extending this approach to other crop species could pave the way for a scalable, automated plant health monitoring system—advancing precision agriculture and reducing crop losses.

In conclusion, the results confirm that lightweight deep learning architectures, when carefully selected and optimized, can achieve high-accuracy disease detection while remaining computationally efficient, offering a viable solution for large-scale agricultural monitoring and smart farming applications.

